# A humanized *Galleria mellonella* model reveals prophage-mediated breakdown of colonization resistance against *Salmonella*

**DOI:** 10.64898/2026.06.24.734175

**Authors:** Zachary M. Bailey, Lavisha Parab, Karina Krammer, Abigail Dustour, Ricardo Léon Sampedro, Mathilde Boumasmoud, Carolin C. Wendling

## Abstract

**Background:** Colonisation resistance provided by the gut microbiota is a critical barrier to pathogen invasion, yet its study in vivo is constrained by the complexity and cost of vertebrate models. Here, we developed a humanised *Galleria mellonella* infection model by inoculating wax moth larvae with complex human faecal microbiota. 16S rRNA gene sequencing confirmed stable, reproducible establishment of a diverse human-associated community across larvae over four days.

**Results:** Humanised larvae exhibited colonisation resistance against *Salmonella enterica* serovar Typhimurium, with mortality reduced to 20% compared to 90% in non-colonised controls. To test whether prophages could overcome this barrier, we infected larvae with isogenic *S*. Tm strains differing in the presence of prophage P22. Infection with the P22-carrying strain resulted in a threefold higher larval mortality (60% vs. 20%), increased pathogen load, and a significant reduction in the abundance of resident *E. coli*. Free P22 virions were detected early after infection, indicating extensive prophage activity. Notably, P22 can neither adsorb nor lyse resident *E. coli,* indicating that prophage-mediated invasion success did not rely on direct lysis. Instead, using high-throughput metabolic profiling paired with whole genome sequencing of three replicate lineages, we found that phage activation intensified resource-partitioning, accelerating functional metabolic adaptations in *E. coli* that significantly reduced the niche-overlap between the invading pathogen and the commensal *E. coli*.

**Conclusion:** Our findings establish the first humanised *G. mellonella* model supporting complex human microbiota and provide a novel non-lytic mechanism by which prophages influence species interactions. This scalable, low-cost model offers a new platform to dissect pathogen–phage–microbiota interactions relevant to human gut ecology.

## INTRODUCTION

Colonisation resistance by the gut microbiota is a fundamental mechanism protecting hosts against infection by enteric pathogens such as *Salmonella enterica* [1]. Disruptions to this barrier, for example, through antibiotics or pathogen-induced microbiome alterations, can increase pathogen colonisation and disease severity [2–4]. While vertebrate models, particularly germ-free or antibiotic-treated mice, have been essential for studying colonisation resistance and pathogen–microbiome interactions [5–8], these models are costly, low-throughput, and experimentally constrained.

The greater wax moth (*Galleria mellonella*) has become a popular invertebrate infection model, offering a simple, scalable, and ethically flexible system to study microbial pathogenesis and host–microbe interactions [9–12]. Larvae can be easily infected and monitored for survival, pathogen load, and immune responses [9–12]. However, their natural gut microbiome is typically low in diversity and dominated by *Enterococcus* spp. [13]. This limits ecological realism and direct studies of colonisation resistance conferred by complex human-associated communities.

Incorporating human-derived microbiota into *G. mellonella* could therefore create a novel, affordable, and high-throughput in vivo model to study microbiome–pathogen interactions. While human-microbiota-associated (HMA) mice are well established [14], the feasibility and utility of such an approach remain largely unexplored in invertebrate systems.

Beyond the composition of the microbiota itself, bacteriophages, especially prophages integrated into pathogen genomes, can play critical roles in shaping infection dynamics. Prophages are widespread across bacterial genomes, including commensal taxa, but comparative genomic studies and large-scale genome mining shows that prophage are more prevalent in pathogens compared to commensals. A simple explanation is that prophages often encode accessory functions associated with pathogenic phenotypes [15–18], including virulence factors and toxins [19–21], functions that facilitate immune evasion [22] or promote biofilm formation [23]. Beyond that, prophages can accelerate bacterial evolution [24,25] and act as self-replicating weapons: upon induction, they lyse susceptible resident bacteria, which reduces competition [26–29]. However, this only works if the released phage can actively infect and kill competitors. Given the extreme specificity of phages, this is rather rare and it leaves a significant gap in our understanding if and how prophages impact non-susceptible species that occupy an overlapping niche.

To answer this question, we developed a human-microbiota-associated (HMA) *G. mellonella* model by colonising larvae with a complex human faecal microbiota. We then tested whether prophages could disrupt microbiome-derived colonisation resistance by infecting larvae with isogenic *Salmonella enterica* serovar Typhimurium strains differing only in the presence or absence of the temperate prophage P22, which can neither adsorb to nor lyse commensal *E. coli* from our faecal communities.

We found that prophage carriage increases pathogen invasion success in vivo, associated with higher larval mortality, increased pathogen load, and reduced growth of resident *E. coli.* As *E. coli* is likely the strongest competitor to *Salmonella* due to their shared ecological niche, we quantified the metabolic niche overlap and found that prophage activation reduces competition by forcing *E. coli* to shift its core metabolic profile. High-throughput phenotypic profiling and whole genome sequencing of parallel replicate lineages revealed that this metabolic shift was driven by treatment-specific mutations.

## RESULTS

### Establishment of a well-tolerated, stable human-microbiota-associated (HMA) *Galleria mellonella* model

We first established a human-microbiota-associated (HMA) *Galleria mellonella* model by colonising larvae with complex human faecal microbiota. 16S rRNA gene sequencing confirmed successful establishment of a diverse bacterial community that persisted at similar levels over four days (Fig 1a). Alpha diversity (Shannon index) was high and stable across time points in uninfected larvae (Fig 1b), and community composition was reproducible across biological replicates. The established community included typical human-associated taxa such as *Bifidobacterium*, *Bacteroides*, *Blautia*, *Lachnoclostridium*, and *Escherichia–Shigella* (Fig. 1a). Sequencing of the original faecal slurry revealed a community dominated by the same human-associated genera that subsequently established in the *G. mellonella*, indicating efficient transfer of the main inoculum members into the larval gut. However, once inside the larval host, *Escherichia*-*Shigella* became comparatively more dominant than in the original faecal slurry (Fig 1a), suggesting that the larval gut environment favours the expansion of facultative anaerobes while still supporting a broad set of human-derived commensals.

**Figure 1:**
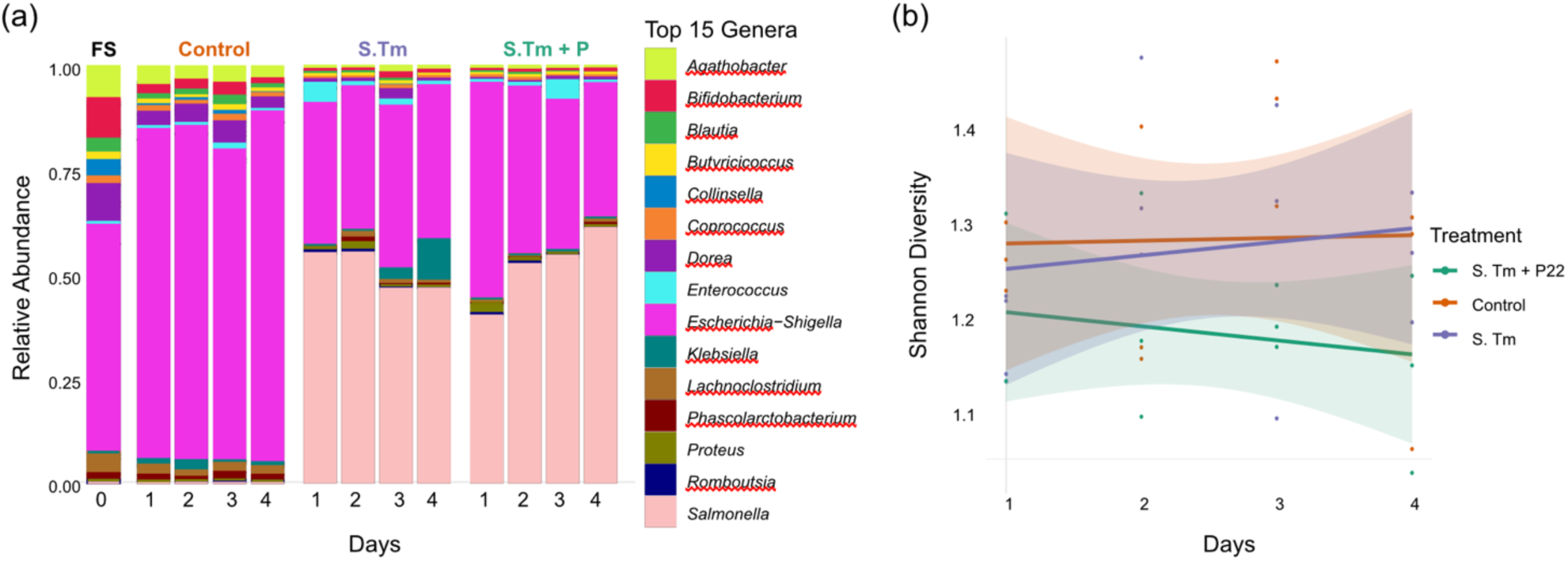
HMA-larvae contain a diverse and stable community composition. **(a) Relative abundance of the top 15 bacterial genera.** Bar plots display taxonomic profiles over a 4-day longitudinal timeline following pathogen invasion. Treatments include the initial fecal sample (FS), uninfected larvae (Control), which only contain the faecal sample but were not infected with *Salmonella* Typhimurium *S*. Tm, larvae infected with prophage-free *Salmonella* Typhimurium (*S*. Tm), and larvae infected with P22 prophage-carrying *Salmonella* Typhimurium (*S*. Tm + P). Taxa are color-coded at the genus level according to the legend. Individual bars represent independent biological replicate pools for each sampling day. Only one replicate out of three is shown. All replicates are shown in Figure S1 **(b) Longitudinal shifts in microbial alpha diversity.** Line trajectories track community complexity measured via Shannon Diversity Index across a 4-day colonization timeline. Colored lines represent linear regression trends for each treatment cohort: uninfected control (orange), *S*. Tm (purple), and *S*. Tm + P22 (green). Shaded area indicates the 95% confidence intervals around the estimated group means. Dots display raw Shannon diversity index metrics for individual biological replicates (n=3).

Colonisation of *G. mellonella* larvae with the human microbiota was well tolerated, with only ∼10% background mortality over the four-day experimental period (Fig. 2a). Compared to the control groups (no inoculation or SM buffer inoculation), we observed no significant difference in mortality (log-rank χ^2^2=0.00086, p = 1.00). This indicates that neither the oral injection procedure itself nor the introduction of complex human microbiota is pathogenic or stressful for the larvae. Together, our data show that oral inoculation with human faecal slurry establishes a well-tolerated, stable, and diverse human-like gut microbiota in *G. mellonella*, in contrast to the native low-diversity community that is typically dominated by *Enterococcus* spp. [13].

**Figure 2.**
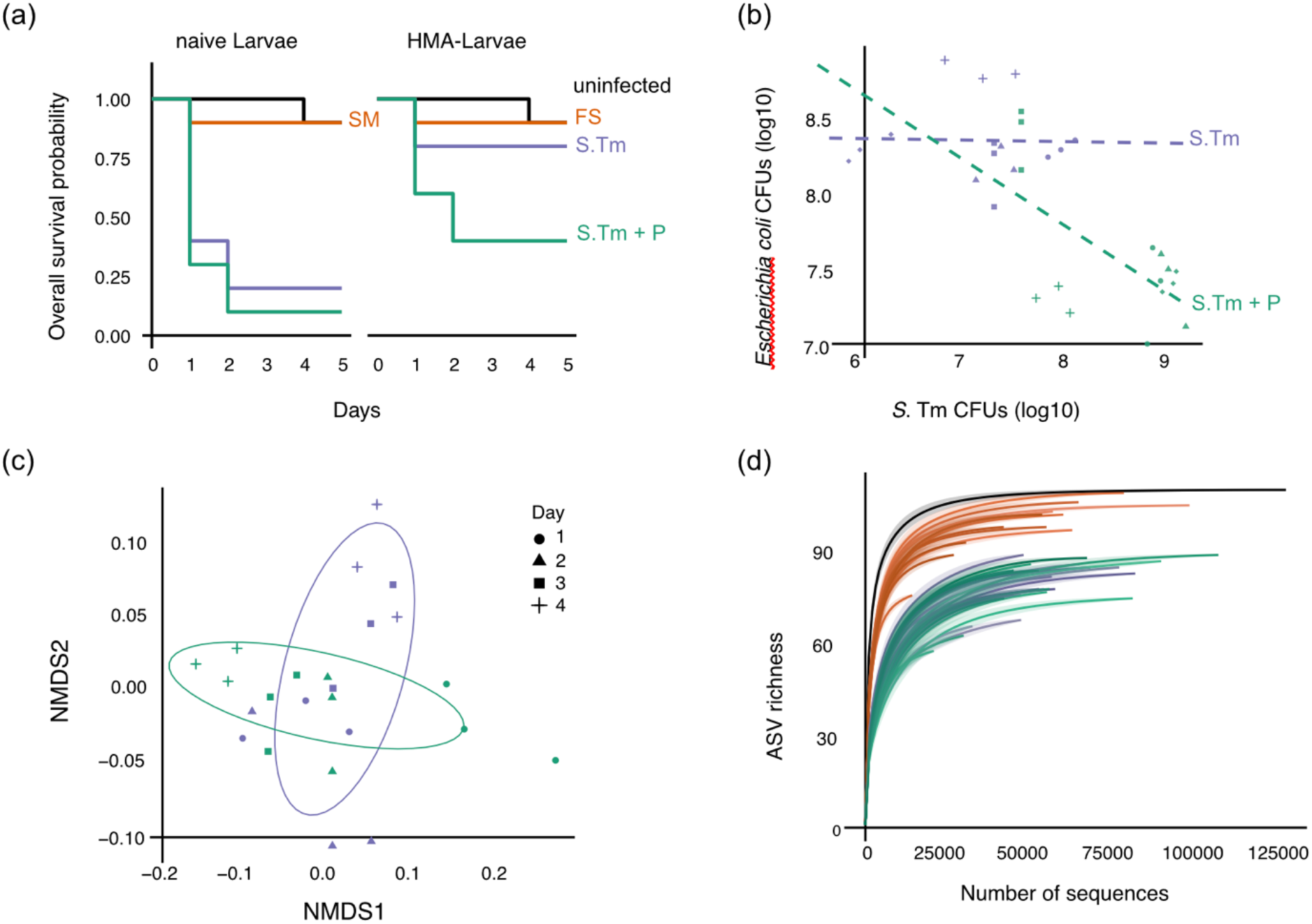
Prophage carriage alters larval survival, competitor colonization dynamics, and intestinal microbiota structure. **(a) Kaplan-Meier survival curves of naive and HMA larvae.** Overall survival probability is tracked across five days, with faecal sample inoculation occurring on Day −1 and *Salmonella* infection occurring on Day 0. Panels compare non-HMA larvae without (left) and human microbiota-associated (HMA) larvae (right). Colors correspond to treatment: uninfected controls (black), basal medium/fecal inoculum controls (FS/SM, orange), prophage-free *Salmonella* Typhimurium (S. Tm, purple), and P22 prophage-carrying *Salmonella* Typhimurium (S. Tm + P, green). **(b) Correlation between *Escherichia coli* and *S.* Tm colonization loads.** Scatter plot displays the relationship between resident *E. coli* abundance (log₁₀ CFU) and *S.* Tm pathogen load (log₁₀ CFU) recovered from larvae homogenates. Dashed lines indicate linear regression trends for the prophage-free track (S.Tm, purple, R^2^ =0.00, p=0.929) and prophage-active track (S.Tm + P, green, R^2^ =0.38, p=0.014). Individual data points represent distinct biological replicates, shapes correspond to independent sampling days: Day 1 (circle), Day 2 (triangle), Day 3 (square), and Day 4 (cross). **(c) Non-metric multidimensional scaling (NMDS) of microbiota beta diversity.** Ordination plot is derived from microbial community distance profiles, with confidence ellipses (95%) color-coded by infection track (S. *Tm*, purple; S. *Tm* + P, green). Shape corresponds to sampling days. **(d) Rarefaction curves of intestinal bacterial communities.** Rarefaction profiles display alpha diversity sample coverage, plotting observed Amplicon Sequence Variant (ASV) richness against total sequencing depth (Number of sequences). Curves are color-coded according to treatment: original faecal slurry (black), uninfected controls with fecal inoculum (orange), *S*. Tm infection (purple), and *S*. Tm + P22 infections (green).

### HMA Larvae exhibit colonization resistance against pathogenic *Salmonella enterica*

To assess whether human microbiota confers colonisation resistance in vivo, we infected HMA larvae and naïve control larvae with *Salmonella enterica* subspecies *enterica* serovar Typhimurium (*S.* Typhimurium) ATCC 14028. HMA larvae showed significantly improved survival. Specifically, mortality in naïve larvae reached up to 80% and 90% upon infection with either *S.* Tm or *S.* Tm + phage P22. In comparison, mortality in response to S. Tm infection in HMA larvae was as low as ∼20% (Log-Rank Test, Chi-Squared = 12.9, p = 0.005 for S. Tm, Chi-Squared = 14.4, p = 0.002 for *S.* Tm +P22; Fig. 2a). These data demonstrate that, like vertebrate infection models, the introduced human microbiota provides functional colonisation resistance against *S.* Tm infection in the *Galleria* model.

### Prophage carriage increases pathogen invasion success in HMA larvae

To test whether prophages can overcome microbiome-derived colonisation resistance, we infected HMA larvae with isogenic *S.* Tm differing only in the presence or absence of the temperate prophage P22. Infection with the P22-carrying strain resulted in a significantly increased pathogen invasion success: larval mortality rose from ∼20% with the P22-free strain to ∼60% with the P22-positive strain, corresponding to a threefold increase in mortality (Log-Rank Test, Chi-Squared = 2.9, p = 0.09; Fig. 2a).

Consistent with the increased mortality, *S.* Tm carrying phage P22 maintained significantly higher CFU/mL loads than phage-free *S.* Tm across days 1–4 post-inoculation (β = 1.38 log₁₀ CFU/mL, F₁,₁₂ = 23.09, p < 0.001, Fig SI), with both treatments showing a modest decline in bacterial load over time (day effect: F₁,₁₂ = 29.18, p < 0.001) and no significant difference in the rate of that decline between treatments (treatment × day interaction: F₁,₁₂ = 1.23, p = 0.289). At 48 h post-infection, mean *S.* Tm CFUs reached 8.9 × 10^8^ in the P22+ group compared to 2.8 × 10^7^ in the P22– group, corresponding to an approximately 30-fold increase (Fig. S2).

Free P22 virions were detected in larval homogenates early after infection, with titres peaking at 6 × 10^2^ PFU/larva at day 1 before declining over subsequent days (Fig. S3). This suggests active prophage induction and release during early infection. These results indicate that prophage carriage and subsequent release can significantly enhance *Salmonella* invasion success in vivo within complex human-derived microbiota.

### Infection and prophage carriage perturb community composition

Infection of HMA larvae with *S*. Tm substantially disturbed the established human-derived gut community. NMDS ordination showed that both infected groups (with and without P22) clustered separately from non-infected controls, indicating marked changes in community structure (Fig. 2c), driven by *S*. Tm. Rarefaction analysis confirmed that infection reduced species richness relative to non-infected larvae (Fig. 2d). This is because *Salmonella* became the dominant genus after infection, thereby displacing commensals such as *Agathobacter*, *Bifidobacterium, Bacteroides, Blautia* and *Escherichia–Shigella* (Fig. 1a).

Alpha diversity (Shannon index) was slightly lower in larvae infected with the P22-carrying strain compared to *S*. Tm alone, although variation across replicates was considerable (Fig. 1b). Importantly, LDA differential abundance analysis revealed distinct patterns between infection groups: while *S*. Tm infection alone enriched for *Proteus* and *Streptomyces*, prophage carriage coincided with additional enrichment of *Salmonella* and *Enterococcus* (Fig. S3). Given that *Enterococcus* is the natural dominant taxon in naïve larvae, this may reflect partial regrowth or niche re-expansion under prophage-mediated disturbance. Together, these data show that *Salmonella* infection itself drives the major community shift and loss of richness, while prophage carriage contributes a modest but distinct effect by altering the abundance of specific taxa.

### Prophage carriage reduces *E. coli* abundance and reduces niche overlap

In HMA larvae infected with the P22-carrying *Salmonella* strain, we observed a significant decline in the abundance of resident *E. coli* compared to infection with the P22-free strain (F₁,₂ = 10.26, p = 0.0328; Fig. S1). Importantly, in the presence of P22, *E. coli* abundance was significantly negatively correlated with *Salmonella* load (Pearson’s Product-Moment Correlation, t = −2.85, p = 0.0137, Fig. 2b), suggesting that higher pathogen loads coincide with reduced competitor abundance. P22 can neither adsorb nor lyse the resident *E. coli* (Supplementary material, note S1). Still, both species occupy a similar ecological niche to *S. Tm* [7,30] and can compete for overlapping resources, especially in conjunction with other microbiota [5,7,30]. We thus specifically examined how prophage carriage affected *E. coli* performance, evolution, and abundance in HMA larvae. To support the assumption of niche overlap, we compared carbon source utilization profiles of ancestral and evolved *E. coli* isolates from the inoculated faecal sample and both *S. Typhimurium* strains using BIOLOG EcoPlates. The metabolic profiles showed substantial overlap (Fig. S5), with a Pianka’s overlap index of 0.87 for both ancestral species comparisons. While niche competition declined over time, prophage carriage altered niche competition by reducing the niche overlap by 5% compared to the non-lysogen (Fig 3a).

**Figure 3.**
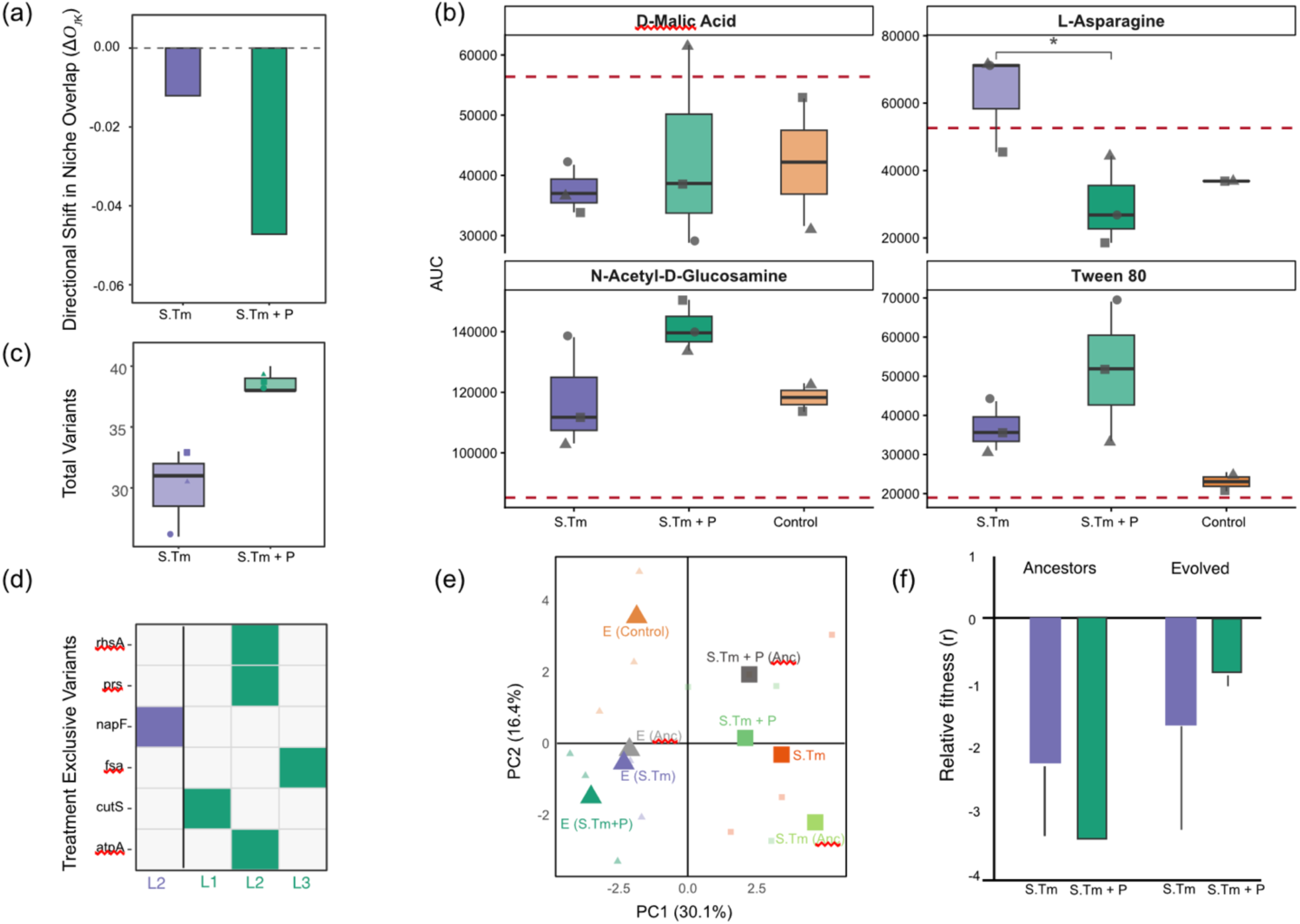
Prophages reduce niche overlap. **(a) Evolutionary shift in dietary niche overlap vectors.** Vertical bars display the directional displacement vector (ΔO_JK_) in Pianka’s niche overlap index over the colonization timeline. Negative values represent character displacement away from the ancestral resource footprint under standard co-evolution (*S*. Tm, purple) and prophage-induced co-evolution (*S*. Tm + P, green). **(b) Mean Growth across four selected substrates.** Integrated curve volumes (Area Under Curve, AUC) across 24-hour incubation for *D-malic acid, L-asparagine, N-acetyl-D-glucosamine,* and *Tween 80*. Individual biological replicate data are shown by different shapes: Lineage 1 (circle), Lineage 2 (triangle), and Lineage 3 (square). Dashed red horizontal lines correspond to ancestor baseline levels. The brackets indicate statistically significant treatment effects based on Tukey’s HSD post-hoc contrasts (*, *p* < 0.05). The four substrates were selected because they had the strongest difference in growth between E. coli evolved in the presence and absence of P22. All substrates are shown as heatmap in Figure S5. **(c) Total variants in evolved E. coli.** Boxplots display the total number of treatment-exclusive nucleotide variants accumulated per sample. Colors correspond to treatment: *S*. Tm (purple), *S*. Tm + P (green). **(d) Treatment specific variants with a high impact on protein function split per lineage.** Shown are treatment-exclusive, non-synonymous mutational variants. Colors correspond to treatment *S*. Tm (purple), *S*. Tm + P (green). Filled squares indicate the presence of independent variants inside metabolic and transport genes (*atpA, cutS, fsa, napF, prs, rhsA*). **(e) Principal component analysis (PCA) of *E. coli* phenotypic profiles.** The ordination plot displays the variation in metabolic profiles based on raw AUC values, with the percentage of variance explained by the first two principal components indicated on the axes (PC1: 30.1%, PC2: 16.4%). Large triangles indicate the group centroid means for each treatment group. Small symbols represent individual biological replicates (n=3). **(f) Relative competitive fitness during pathogen invasion.** Bar charts visualize relative fitness coefficients (*r*) of ancestral (left) and evolved (right) *E. coli* derived from direct pairwise assays against *S.* Tm (purple) or *S.* Tm+P22 (green). Shown are means ± s.e. (n=3).

To identify the specific metabolic drivers behind these metabolic shifts, we focused on substrates where we observed the biggest difference in AUC between *E. coli* evolved in the presence or absence of P22. Isolates from the phage treatment exhibited a strong decline on key amino acids, specifically L-asparagin and L-serine, combined with a distinct gain of growth D-malic acid, N-acetyl-D-glucosamine, and Tween 80 (Fig. 3b).

To uncover the genetic mechanisms underlying these metabolic shifts, we sequenced evolved *E. coli* from all replicate lineages (n=3). After removing one sample that identified as *Enterococcus faecalis*, comparative genomic analysis against the ancestral isolate revealed treatment-specific mutations. In the presence of P22, *E. coli* accumulated significantly more variants compared to P22-free treatments (Mann-Whitney test, W=9, p = 0.038, Fig. 3c), whereby all changes occurred inside functional pathways.

Instead of relying on a single parallel mutational pathway, different lineages acquired different combinations of mutations, which altered their capacity on separate substrates (Fig. 3d). This multi-route genetic divergence is directly reflected in the phenotypic distribution of individual replicate lineages (Fig. 3b). For instance, lineage 1 acquired a single mutation in the envelope-associated sensory kinase cutS. In contrast, lineage 2 acquired a unique combination of mutations in atpA (proton motive force regulation), prs, and rhsA. This specific alteration in proton gradient energetics and sugar synthesis can explain why lineage 2 is visible as a phenotypic outlier that drives the observed reduction in growth on L-asparagine coupled with the highest growth on D-malic acid (Fig 3b). Lastly, lineage 3 carried a treatment-exclusive variant in fsa (fructose-6-phosphate aldolase), which could explain its high growth on N-acetyl-D-glucosamine (Fig 3b). While these individual lineages relied on entirely different genetic outcomes, their phenotypic trajectories ultimately converged along the same evolutionary trajectory determined by standard co-evolution with *Salmonella* (Fig. 3e). This demonstrates that the presence of the prophage P22 accelerated the existing co-evolutionary pressures rather than forcing a completely new phenotypic path.

To infer the ecological consequences of this reduced niche-overlap, we performed direct competition assays between ancestral or evolved *E. coli* clones and isogenic *S*. Tm strains. Across all conditions, *S*. Tm outcompeted *E. coli*, consistent with its role as an invasive pathogen. However, the magnitude of this competitive disadvantage differed: when competing against the P22-carrying *Salmonella* strain, evolved *E. coli* showed a smaller fitness gap compared to ancestral clones (Fig. 3f). By contrast, when competing against the P22-free *Salmonella* strain, there was no difference in net fitness between ancestral and evolved *E. coli*. This indicates that by shifting its metabolic profile away from Salmonella’s, the commensal narrows its competitive disadvantage. While the prophage initially drives pathogen invasion success, it ultimately increases the competitive fitness of the commensal, suggesting that non-lytic prophage-mediated invasion success is transient and counterbalanced over time by rapid evolutionary responses in the microbiota.

## CONCLUSION

By establishing an HMA *G. mellonella* model, we introduce a scalable platform that bridges the gap between high-throughput in vitro models and low-throughput vertebrate systems. Combining functional and mechanistic tests, our study uncovers a previously underappreciated ecological role for prophages in pathogens: prophage carriage enhances pathogen invasion success in vivo by transiently outcompeting key competitors. However, this effect is short-lived as metabolic adaptations rapidly evolve within the resident competitors.

With that, our study complements recent observations in gnotobiotic mice where lytic phages targeting protective commensals were shown to undermine colonization resistance and enable *Salmonella* invasion [31]. By extending this principle from lytic phage-mediated invasion to prophage release triggered by an invading pathogen, we highlight a broader ecological role of prophages in pathogens — one that extends beyond their well-recognized contributions to virulence factor and ARG carriage and instead positions prophages as active mediators of microbial competition, colonization, and community restructuring [26,32–34]. While we observed a massive viral burst peaking at 24 hours post-infection, the exact molecular triggers for this induction and the precise role of these free virions remain unknown. However, their rapid release suggests a massive viral disruption in the gut landscape during the early stages of infection, which accelerated metabolic adaptation five-fold compared to competition with P22-free Salmonella. However, multiple genetic options seem to be possible rather than a single mutational pathway to adapt to P22-triggered stress. These alternative evolutionary routes reshape the commensal’s metabolic profile, narrow the niche overlap by 5% and reduce the commensal’s competitive disadvantage.

This is another example that demonstrates the capacity of prophages to shape community composition and invasion dynamics in densely populated ecosystems, such as the mammalian gut, with cascading ecological and health impacts. Importantly, the transient nature of this advantage helps explain why pathogenic bacteria tend to harbour more prophages than commensals [35]: even a short-lived competitive advantage during invasion may be strongly favoured in the evolution of enteric pathogens.

## METHODS

### 1. Ethics Statement

Stool samples for this study were from consenting anonymous human donors. The sampling protocol was approved by the ETHZ ethics commission: EK2020-N150.

### 2. Study System

We obtained the temperate phage P22 DSM18523 from the DSMZ microbial collection. Infection experiments were performed with the pathogen *Salmonella enterica* subspecies *enterica* serovar Typhimurium (S. Typhimurium) ATCC 14028 containing a chromosomal antibiotic resistance marker (marT:cat). The strain was originally obtained from Wolf-Dietrich Hardt [33,36]. we created *S*. Tm. lysogens containing P22 integrated into the bacterial genome and confirmed lysogeny through PCR using the forward primer 5’-ATGGTGGCAGGAGTTAATGC – 3’ and the reverse primer 5’-CAAACAAATCCCGAACGACT – 3’ covering the sieB (superinfection exclusion B) gene. Unless otherwise mentioned, strains were grown in LB at 37° C at 180 RPM.

We conducted the in vivo infection experiments on last instar larvae of the greater wax moth, *Galleria mellonella* supplied by Baitexpress AG, Switzerland. We stored larvae at 4-8° C after receiving them. Larvae are separated into petri dishes before infection, and only cream-coloured, uninfected larvae were used for our infection experiments. We inoculated the larvae with human microbiome-like communities, following the procedure described below.

### 3. Stool Samples

Stool samples were collected and processed according to Baumgartner et al. (2020), with some modifications [37]. Samples were collected at the Department of Environmental Systems Science, ETH Zürich, on 24th May 2022. Briefly, inclusion criteria for individuals over 18 years old were: non-obese, not in post-surgery recovery, no antibiotics, and no SARS-CoV2 infection within the past six months. Each sample was collected in a 500-ml plastic specimen container (Sigma-Aldrich) and kept anaerobic using an AnaeroGen anaerobic sachet (Thermo Scientific, Basel, Switzerland). We randomly selected one sample from a larger number of donated samples for the experiment. Stool samples were collected in the morning before the experiment and kept for a maximum of 1 h before processing as described in Baumgartner et al. (2020) [37].

To prepare a faecal slurry from each sample, we resuspended 20 g of the sample in 200 ml anaerobic peptone wash (Sigma-Aldrich) containing 1 g/l peptone, 0.5 g/l L-Cysteine, 0.5 g/l bile salts, and 0.001 g/l Resazurin. This resulted in a 10% (w/v) faecal slurry. We then stirred the slurry for 15 min on a magnetic stirrer for homogenization, followed by 10 min of resting to allow sedimentation. From this mixture, we took 100 µl to initiate a human gut microbiome-like culture, thereby enriching the bacterial populations present. For each sample, we used ∼7 ml of basal medium samples (Sigma-Aldrich) containing 2 g/l Peptone, 2 g/l Tryptone, 2 g/l Yeast extract, 0.1 g/l NaCl, 0.04 g K2HPO4, 0.04 g/l KH2PO4, 0.01 g/l MgSO4×7H2O, 0.01 g/l CaCl2×6H2O, 2 g/l NaHCO3, 2 ml Tween 80, 0.005 g/l Hemin, 0.5 g/l L-Cysteine, 0.5 g/ l bile salts, 2g/l Starch, 1.5 g/l casein, 0.001g/l Resazurin. The pH was adjusted to 7, and 0.001g/l Menadion was added after autoclaving. Samples were grown at 37° C in a Coy Model A Vinyl Anaerobic Chamber to enrich the microbiome and subsequently stored at −80° C with 25% Glycerol.

### 4. Community inoculation

To inoculate the larvae with human microbiome-like communities, we diluted each enriched faecal sample to a final concentration of 10^4^ CFU/ml of culturable bacteria in Sodium-Magnesium (SM) buffer (5.8 g NaCl, 2 g MgSO4 7 H2O, 50 ml 1M Tris HCl, pH 7.5 for 1 L of buffer). Importantly, we note that culturable bacterial strains are only a subset of the total composition of the human gut microbiome present in culture [38,39]. Subsequently, we orally injected 5 µl of this culture between the mandibles of *G. mellonella* larvae using a Hamilton glass syringe and 27-gauge Sterican needle, following the oral injection method as described in Coates et al. and Harding et. al. [11,40]. To account for the potential impacts of the faecal sample itself, we injected a group of larvae with SM buffer alone as a control. Additionally, to control any harm caused during handling, we monitored a naive group of larvae during each infection experiment.

### 5. Characterization of Starting Communities

#### a. Microbial Community Composition

We plated the faecal samples on chromatic agar, chromatic agar with chloramphenicol and Hektoen enteric agar plates grown at 37° C. We observed two species, i.e., *Escherichia coli* and *Enterococcus spp*., that were able to grow on the chromatic agar and Hektoen enteric agar plates. Humanized larvae samples contained both *Escherichia coli* and *Enterococcus spp.* determined by plating on chromatic agar plates.

#### b. Clonality of the dominant *E. coli*

We isolated six *Escherichia coli* colonies from FS10 and conducted a repetitive element palindromic PCR (rep-PCR) using the primers REP1R (5’-NNNGCGCCGNCATCAGGC-3’) and REP2 (5’-ACGTCTTATCAGGCCTAC-3’) [ https://doi.org/10.1093/nar/19.24.6823]. PCR conditions were 94° C for 4 min, 30 cycles of 94° C for 1 min, 40° C for 1 min, 65° C for 8 min, and a final 65° C for 16 min. Amplified DNA was analysed by gel electrophoresis (2% agarose, 130V, 1h) to compare strain-specific profiles. We also whole-genome sequenced (Oxford Nanopore Technologies, Plasmidsaurus) the clones and determined the Average Nucleotide Identity (ANI) of the genomes using FastANI v1.34 [https://doi.org/10.1038/s41467-018-07641-9] with standard settings. This step confirmed that all randomly selected E. coli from this faecal sample had an ANI of 99.99% and can thus be considered as clonal.

#### c. Phage Adsorption and Susceptibility testing

To determine the susceptibility of the dominant strains of the culturable bacteria in each community, *E. coli* and *Enterococcus*, to phage P22 or the native prophages of the focal pathogen Gifsy-1, Gifsy-2, Gifsy-3 and ST64B, we conducted adsorption and standard spot-assays [41,42]. We spotted the lysates of P22-containing *S*. Tm and P22-free *S*. Tm onto lawns of six randomly selected *E. coli* and *Enterococcus* clones derived from the faecal sample. No plaque formation indicated the resistance of the clone to phage P22 or any of the native prophages of *S*. Tm.

We measured adsorption of P22 to six ancestral *E. coli* clones, with the P22-free *S*. Tm as a positive control. We mixed ∼10⁸ exponentially grown bacteria with P22 lysates (∼10⁵) and incubated the mix for 20 minutes at 37 °C, 180 rpm. We sampled 500 μL after 10 and 20 minutes, filtered it through 0.22 μm filters, and enumerated the PFUs to measure adsorption in this time interval. To account for sampling and pipetting errors, we measured PFUs using 2-3 technical replicates.

#### d. Inhibition Assay

We diluted the processed faecal sample 1:1 with PBS and centrifuged it for 5 minutes at 4696 RCF. We spotted the supernatant of the faecal sample on the *S*. Tm to see if any of them could inhibit *Salmonella* growth. We made overlay LB agar plates and mixed 3ml of soft agar with 100 µl of *Salmonella* cells suspended as a concentrated slurry (∼10^10^ CFUs/ml) in 10 mM MgSO4.

### 6. Infection Experiment

To test the effect of prophages on a complex community, we infected 45 larvae with a random faecal sample, following our community inoculation protocol. We followed up our community inoculation the next day, infecting 15 larvae with *S.* Tm and 15 larvae with *S.* Tm + P22. We kept the remaining 15 larvae as uninfected control. After infection, we monitored changes in the population size of the pathogen, free P22 virions, and the culturable members of the gut microbial community, i.e., *E. coli* and *Enterococcus*. To do so, we sacrificed three randomly selected larvae from each group per day. Each sacrificed larva was surface sterilized with 70% ethanol, placed in a 2ml Eppendorf tube, and homogenized for 5 minutes using a Retsch MM301 Mixer Mill Pulverizer. We then mixed 1ml of SM buffer with the homogenate, vortexed the sample for 10 seconds, and performed serial dilutions of the homogenate in SM buffer placing 20 µl in 180 µl of SM buffer. We plated 100 µl of the homogenate solution diluted 10^-5^ onto chromatic agar plates that allow visual discrimination of different taxa. Additionally, we plated another 100 µl of diluted homogenate onto Hektoen enteric agar plates supplemented with 25 µg/ml Chloramphenicol to select for *S.* Tm. To quantify free P22 virions, we mixed 100 µl homogenate with 10 µl of Chloroform and vortexed followed by centrifugation of 4,000 g for 5 minutes in an Eppendorf 5425R Centrifuge. We then serial diluted in SM buffer and spotted this lysate onto overlay plates containing a P22 susceptible *S.* Tm strain performing a standard spot assay [41,42].

In parallel to the 45 larvae described above, we inoculated another 30 larvae with the same randomly selected faecal sample and another 30 larvae with SM buffer. After infection, we allowed the larvae to acclimate to room temperature for 1 hour before incubating at 37° C. We allowed the community 24 hours to establish itself, before infecting 10 larvae from each batch with *S.* Tm and another 10 larvae from each batch with P22-carrying *S.* Tm. We used an inoculum size of 1000 ± 500 CFU and monitored larval survival for five days. A larva was considered dead if it was unable to right itself after being flipped onto its back.

### 7. Whole genome sequencing

Whole genomic DNA was extracted from randomly selected clones of *Salmonella enterica* serovar Typhimurium (S. Tm), and *Escherichia coli.* We selected one ancestral clone of each *Salmonella* and three clones of each evolved *E. coli* and *Salmonella* from the final day of each treatment using a GenElute Bacterial Genomic DNA Kit (Sigma-Aldrich). Libraries were sequenced using PacBio HiFi long-read sequencing at the Functional Genomics Center Zürich. All genomes were initially processed through the *StrainCascade* pipeline [43], which performs assembly and primary quality control. The Flye-based assemblies produced by this pipeline and raw HiFi FASTQ files were used as the starting point for all downstream analyses.

Three samples were identified as containing reads originating from both *S.* Tm and *E. coli*. For these contaminated samples, the *Flye* assembly contigs were classified by aligning them to two reference genomes, the ancestral *S.* Tm genome and a complete *E. coli* K-12 assembly, using minimap2 v2.20 [44]. For each contig, alignment identity and query coverage were computed from the resulting PAF files, and each contig was assigned to the species yielding the higher combined alignment score. Species-specific reads were then extracted by mapping the original HiFi reads back to the classified contig sets using minimap2 with the map-hifi preset, filtering for mapped reads with *SAMtools* [45], and converting the output to FASTQ format. De novo re-assembly of the separated read sets was performed using *Flye* v2.9 with the --pacbio-hifi flag, using genome size estimates of approximately 4.7 Mb for *E. coli* and 4.9 Mb for *S.* Tm [46].

All final genome assemblies were annotated using *Bakta* [47] and screened for prophage content using *PHASTEST* [48]. Single-nucleotide polymorphism (SNP) and indel variant calling was performed using Snippy v4.6 in single-end mode (--se). [49], comparing the *E. coli* and *S.* Tm isolate genomes to their respective annotated ancestral reference genomes in GenBank format (.gbk). Within each Snippy run, reads were aligned to the relevant reference genome using BWA [50], variants were called using *FreeBayes* [51], and predicted functional effects of each variant were annotated using SnpEff [52]. For *S*. Tm isolates, two distinct reference genomes were used to reflect the experimental design: isolates from the SP treatment group (Salmonella co-cultured with P22 phage; FS10_R1/R2/R3_SP) were compared against the annotated assembly of PE_036_P22, a P22 lysogen of the ancestral *S*. Tm strain, whilst isolates from the S treatment group (Salmonella without phage; FS10_R1/R2/R3_S) were compared against the annotated assembly of PE_036, the non-lysogenised ancestral strain; this distinction ensured that prophage-associated genomic differences were not erroneously called as experimental variants. For *E. coli* K-12 isolates, all samples were compared against the highest-quality ancestral *E. coli* assembly, selected automatically on the basis of N50 and contig count from six candidate reference assemblies. Three samples identified as containing mixed *S*. Tm and *E. coli* reads (FS10_R1_SP_E, FS10_R2_S_E, and FS10_R3_SP_E) were first subjected to taxonomic read classification using *Kraken2* (database: k2_standard_16gb_20240605) [53], with species-specific reads extracted using *KrakenTools* [54] prior to Snippy analysis; extracted reads from these samples were additionally renamed to avoid read identifier length conflicts with BWA’s SAM format limitations. Results for each sample were written to individual output directories named by barcode identifier, and variant calling outputs were subsequently analysed in R [55] using the *tidyverse* package suite [56].

### 8. Metagenomic Analysis

Two hundred microlitres of *Galleria mellonella* homogenate per sample were sent on dry ice to Microsynth AG (Balgach, Switzerland) for DNA extraction and amplicon sequencing. The V4 hypervariable region of the 16S rRNA gene was sequenced on an Illumina MiSeq platform using paired-end Illumina sequencing, yielding an average of 63,297 reads per sample.

Downstream analysis was performed in R. Amplicon sequence variants (ASVs) were inferred and denoised using the *DADA2* package [57], with sequence manipulation performed using *Biostrings* [58]. Taxonomy was assigned against the SILVA 16S rRNA database version 138.1 [59]. The resulting ASV count table, taxonomic assignments, and sample metadata were integrated into a *phyloseq* object for all downstream analyses [60]. Samples corresponding to the uninfected control group were removed prior to comparative analyses, retaining only samples from the *Salmonella*-infected and P22 lysogen treatment groups.

Alpha diversity and rarefaction curves were assessed using the *ranacapa* package [61]. Community dissimilarity between samples was quantified using Bray–Curtis distances computed with the vegan package [62,63]. The top 20 most abundant genera were visualised as stacked bar charts of relative abundance.

Differential abundance analysis was performed at the genus level. The phyloseq object was agglomerated to genus rank and converted to a *DESeq2* object [64]. Size factor estimation and negative binomial modelling were carried out with *DESeq2* [64], using the “none” (uninfected) group as the reference level. Pairwise contrasts were computed between the Salmonella-infected group versus uninfected controls, and between the P22 lysogen group versus uninfected controls, extracting log∼2∼ fold changes and adjusted p-values (Benjamini–Hochberg correction). Results were visualised as bubble plots in which dot size represents the magnitude of the log∼2∼ fold change and colour indicate the direction of change. Additionally, Linear Discriminant Analysis Effect Size (LEfSe) was performed using both the *microbiomeMarker* [65] and *lefser* [66] packages to identify genera discriminating between treatment groups, with pairwise comparisons conducted to satisfy the dichotomous class requirement of *lefser*.

### 9. Metabolic profiling

We tested the metabolic niche overlap of E. coli and S. tm using BIOLOG EcoPlates™ (BIOLOG™ Inc., California, U.S.A., Catalog No. 1506) which contain 31 carbon sources. We added 99 μl of PBS to all wells followed by mixing and inoculation of 1 μl bacterial overnight culture grown in LB. We incubated the plates for 24 hours at 37° C with shaking at ∼180 rpm in a plate reader (TECAN, Infinite® 200 PRO) and recorded the optical density at 600 nm every 15 minutes.

### 10. Competition assays

To determine differences in fitness between ancestral E. coli and ancestral Salmonella compared to evolved E. coli and evolved Salmonella, we performed competition assays as described in Wendling et. al. (2022) [67]. Briefly, each competition culture was done in triplicates. Overnight cultures of competing strains were mixed 1:1 and 60 µl of this mixture was inoculated to 6 ml LB to initiate each competitive culture. After 24 hours, fitness was estimated by means of plating on ChromAgar. Competitive fitness was estimated as the ratio in Malthusian parameters [68]:

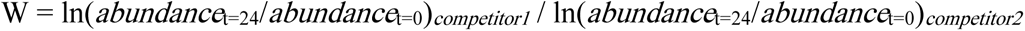

### 11. Statistical analysis

#### a. Infection experiment dynamics

Bacterial and bacteriophage densities were enumerated as colony-forming units (CFU) and plaque-forming units (PFU), respectively. Raw counts were converted to densities per millilitre by multiplying by the relevant dilution factor and were log₁₀-transformed prior to analysis in R [55]; samples with no detectable counts were assigned a value of zero on the log₁₀ scale. For each treatment, sampling day and, where applicable, bacterial taxon (*Salmonella enterica*, *Escherichia coli*, *Enterococcus* spp.), mean log₁₀ densities were summarised together with their standard deviation and standard error (SE = SD/√n, n = 3 replicates) using the *plyr* package [56]. CFU and PFU dynamics over time were visualised as mean ± SE using *ggplot2* [56]. To test whether *S.* Tm densities differed through time between the phage-free and P22-lysogen treatments, a linear mixed-effects model was fitted with log₁₀ CFU/ml as the response, treatment, day and their interaction as fixed effects and replicate as a random intercept, using *lme4* [69], with degrees of freedom and *P*-values obtained via Satterthwaite’s approximation in *lmerTest* [70]. The association between the two taxa was quantified separately for each *Salmonella* treatment (P22-lysogen versus phage-free) using Pearson’s correlation coefficient. Ordinary least-squares regression lines of *E. coli* on *S. enterica* log₁₀ CFU/ml using lm() from *stats* were then fitted within each treatment and overlaid on scatter plots of the paired per-replicate counts to visualise the direction and strength of the relationship.

#### b. Survival analysis

Results from our survival curves were analyzed in R using the Survival of *Galleria mellonella* larvae was analysed in R (version 4.6.0; [55]). Larvae were monitored daily and scored as a binary status (alive or dead), with individuals surviving to the end of the experiment right-censored at the final observation day; right-censored time-to-event objects were constructed with the Surv() function of the *survival* package [71]. Treatment-specific survival probabilities were estimated using the non-parametric Kaplan–Meier estimator from the survival package [71], and differences in survival between treatments were tested using the log-rank test using the survdiff() function of the same package [71], applied both across the full set of treatments and to predefined pairwise subsets of interest (e.g. *Salmonella* lysogen versus *Salmonella*-plus-phage comparisons within each faecal-microbiome background); pairwise comparisons were not adjusted for multiple testing. Kaplan–Meier survival curves were visualised using the *ggsurvfit* [72] and *survminer* [73] packages, built on *ggplot2* [74].

#### a. BIOLOG EcoPlate analysis

Raw kinetic growth data from BIOLOG EcoPlates were processed in R. We excluded wells without growth by applying an empirical growth threshold calculated from the mean plus two standard deviations of the un-evolved ancestral water control wells. For the remaining wells, we integrated the area under the growth curve (AUC) across 24 hours using the trapezoidal rule [75].

To quantify the resource competition and metabolic similarity between *E. coli* and *S*. Tm., we calculated the community niche overlap index using Pianka’s symmetric overlap equation [76]. The index ranges from 0 (completely distinct niches) to 1 (complete niche overlap).

To highlight the most important phenotypic changes, raw delta-yields were calculated by subtracting the mean AUC of the phage-free lineages from the phage-treated lineages. Substrates were ranked hierarchically by the absolute magnitude of this delta.

To infer compounds with statistically significant growth between evolution in the absence and presence of P22, we used a Linear Mixed-Effects Models (LMM) using the *lme4* package [69] with evolution as a fixed effect and independent lineages integrated as a random intercept block to account for batch and flask-specific noise. Pairwise post-hoc comparisons were calculated using Tukey’s honest significant difference (HSD) test via the *emmeans* package [77].

To visualise global phenotypic shifts, principal component analysis (PCA) was performed on the growth-validated AUC matrix using the prcomp function with scaling. Trajectory lines were traced by calculating group centroid means to measure the direction and acceleration of evolutionary selection vectors.

## Notes

### Competing Interest Statement

The authors have declared no competing interest.

### Funding

- SNF: PZ00P3_179743

