## Supplementary Information for "A humanized *Galleria mellonella* model reveals prophage-mediated breakdown of colonization resistance against *Salmonella*"

SUPPLEMENTARY MATERIAL

Note 1: **The O-antigen type targeted by P22 differs from that predicted for the ancestral or evolved *E. coli***

P22 mediates genome injection into its hosts by interactions with the LPS, O-antigen and OmpA of *S.* Tm. All ancestral and evolved *E. coli* clones possessed the O59 O-antigen genes [1] with only minor LPS-core and O-antigen differences. The O-antigen differences are likely not a result of adapting to infection or adsorption by P22 as the O59 O-antigen cannot be targeted by P22. O59 is a GlcNAc-initiated branched antigen with three mannose residues, [α-d-GalA]-α-d-mannose-(1→3)-α-d-mannose-(1→3)-β-d-mannose-(1→3)-α-d-GlcNAc- [2], while the P22 endorhamnosidase specifically targets the galactose-initiated branched antigen [dideoxy-hexose]α-d-mannose-(1→4)-α-l-rhamnose-(1→3)-α-d-galactose-(1→2) [3]. Further, we also tested P22 lysates with different ancestral *E. coli* clones, and P22 neither lyses nor adsorbs to ancestral *E. coli*.

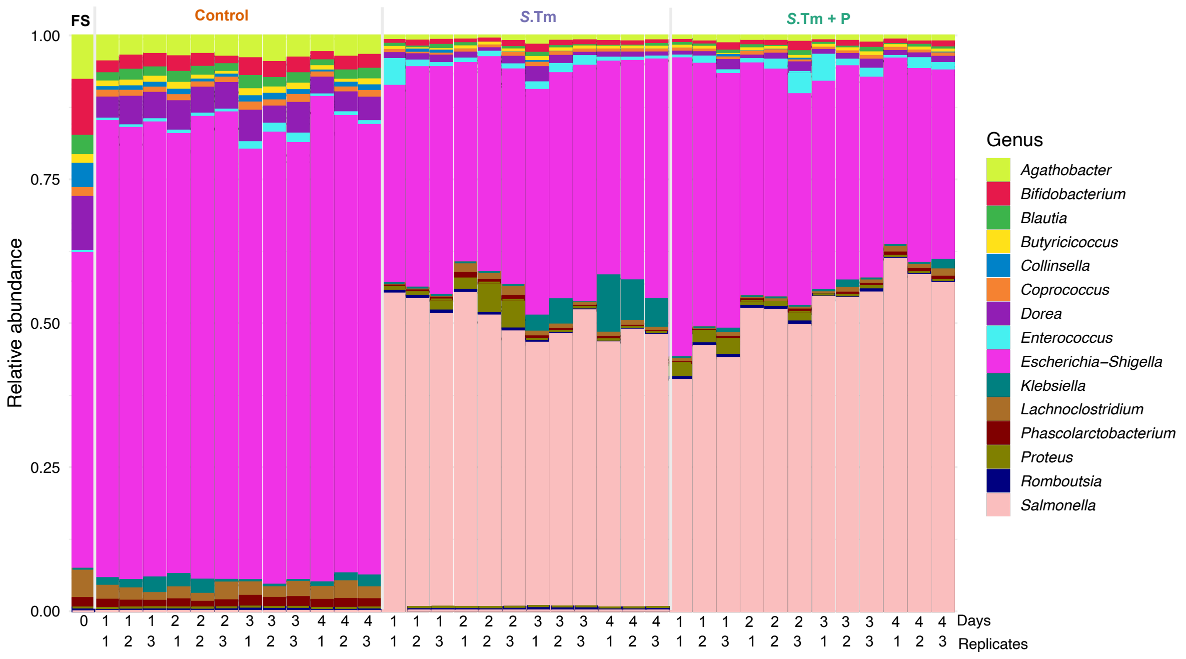


**Figure S1 HMA-larvae contain a diverse and stable community composition**. **(a) Relative abundance of the top 15 bacterial genera shown for all three replicates.** Bar plots display taxonomic profiles over four days. Treatments include the initial fecal sample (FS) that we inoculated into the larvae, uninfected larvae (Control), which only contain the faecal sample but were not infected with *Salmonella*, larvae infected with *S*. Tm, and larvae infected with *S*. Tm + P. Taxa are color-coded at the genus level according to the legend. Individual bars represent independent biological replicates for each sampling day.

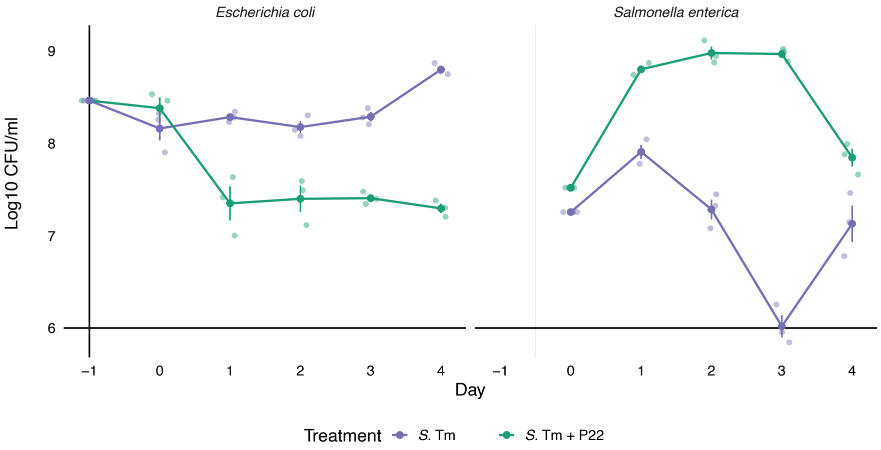


**Figure S2** Mean number of *E. coli* (left) and *S*. Tm (right) quantified as Colony Forming Units (CFU) for each treatment group (*S*. Tm in purple and *S*. Tm + P in green). Shown are means ± s.e. (n=3). Day -1 corresponds to 24 hours past inoculation with faecal sample, Day 0 corresponds to 24h past *Salmonella* infection.

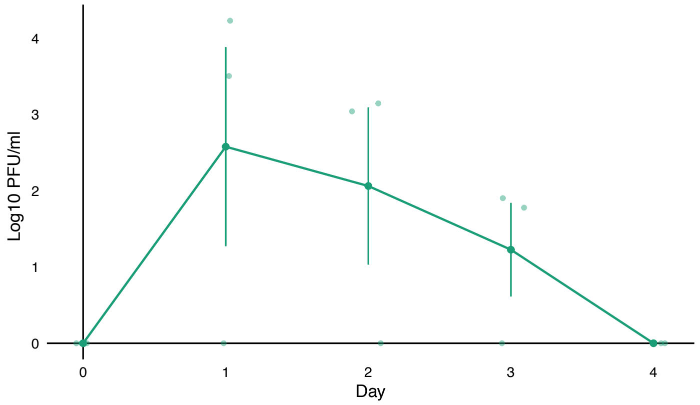


**Figure 3**: Mean number of plaque-forming units (PFU) of phage P22 present in the homogenate of *Galleria mellonella* larvae colonized with fecal sample and infected with *S.* Typhimurium that contains an active P22 prophage. Shown are means ± s.e. (n=3). Day -1 corresponds to 24 hours past inoculation with faecal sample, Day 0 corresponds to 24h past *Salmonella* infection.

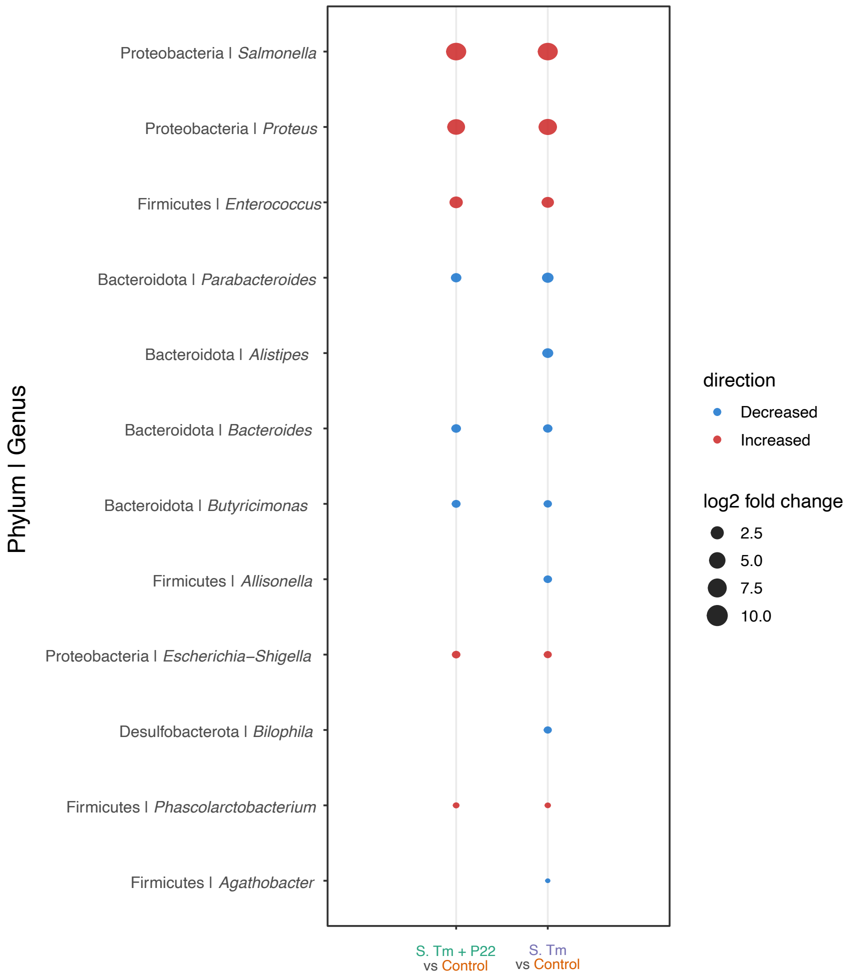


**Figure 4 Differential abundance of bacterial genera based on 16S rRNA gene sequencing**. Genera shown were identified as discriminative by LEfSe (linear discriminant analysis effect size; LDA score > 2.0, $p < 0.05$) for each infection treatment, *S*. Tm, or *S*. Tm + P, relative to the control inoculated with faecal sample only. For each discriminative genus, point size is scaled to the magnitude of the log2 fold change in relative abundance versus the control. Colours correspond to the direction of change (orange increased; blue, decreased). Genera are grouped by phylum (Phylum | Genus).

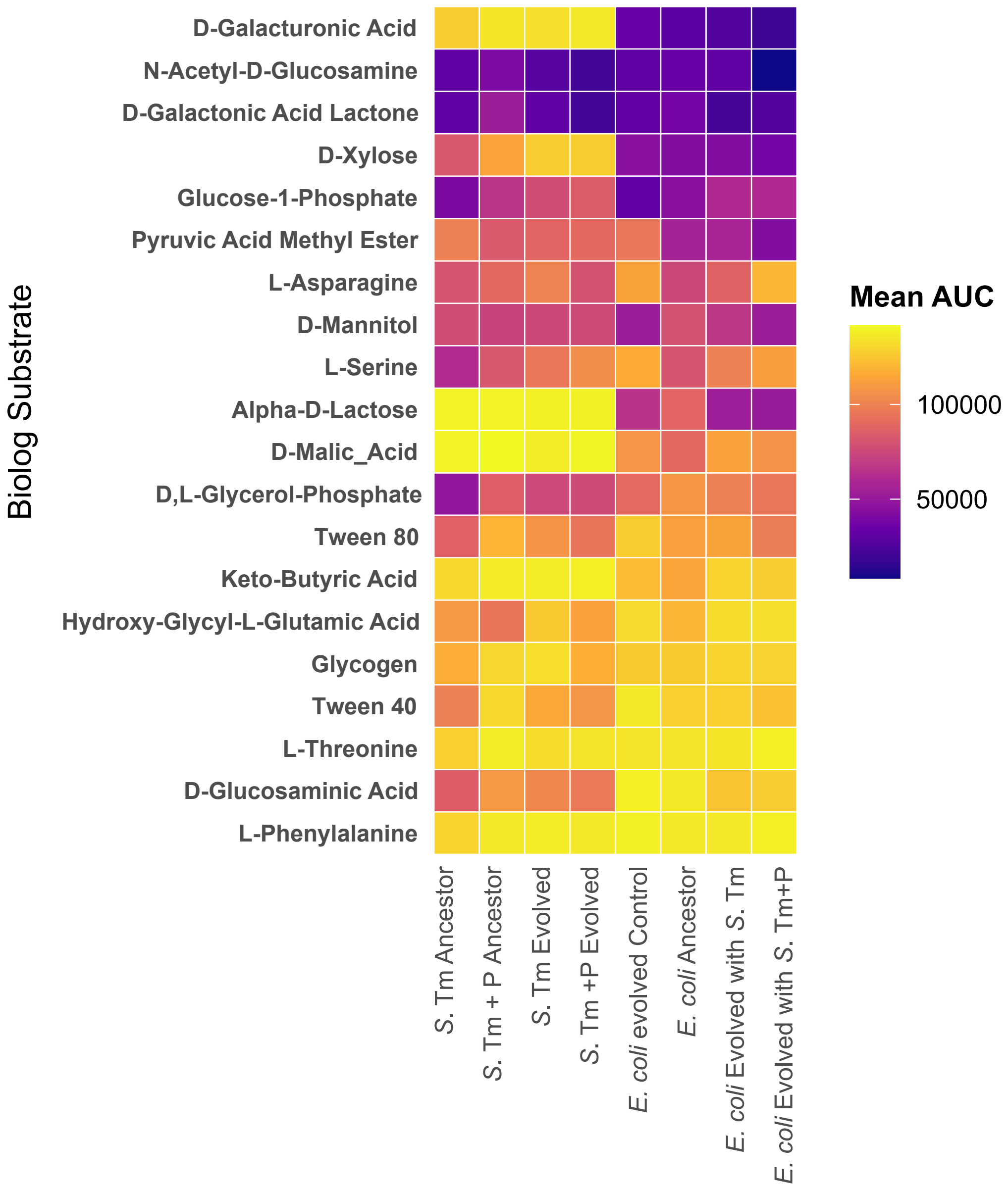


**Figure 5** Growth of ancestral and evolved *S*. Tm, *S*. Tm + P, and *E. coli* on single carbon-source substrates in Biolog EcoPlates. Colors correspond to the Area Under the Curve calculated from 24h OD600 measurements for three replicates per strain (column) on each single carbon source environment (rows); see scale at right. Carbon source environments, where we observed no growth across all isolates are not shown.
